# Vimentin intermediate filaments provide structural stability to the mammalian Golgi apparatus

**DOI:** 10.1101/2022.08.25.505293

**Authors:** Teresa Vitali, Tomasz M. Witkos, Marie F.A. Cutiongco, Guanhua Yan, Alexander A. Mironov, Joe Swift, Martin Lowe

**Author notes:** Present address: Department of Analytical Sciences, AstraZeneca, Cambridge, UK. Present address: School of Chemical and Biomolecular Engineering, Nanyang Technological University, Singapore (Marie F.A. Cutiongco). Summary statement: In this study we uncover a role for vimentin intermediate filaments in providing physical stability to the Golgi apparatus in mammalian cells.

## Abstract

The Golgi apparatus comprises a connected ribbon of stacked cisternal membranes localized to the perinuclear region of most vertebrate cells. The position and morphology of this organelle depends upon interactions with microtubules and the actin cytoskeleton. In contrast, we know relatively little about the relationship of the Golgi apparatus with intermediate filaments. In this study we show that the Golgi is in close physical proximity to vimentin intermediate filaments (IFs) in cultured mouse and human cells. We also show that the *trans*-Golgi network coiled-coil protein GORAB can physically associate with IFs. Although loss of vimentin and/or GORAB does not have major effects upon Golgi morphology at steady-state, the Golgi undergoes more rapid disassembly upon chemical disruption with the drug brefeldin A, and slower reassembly upon drug washout, in vimentin knockout cells. Moreover, loss of vimentin causes reduced Golgi ribbon integrity when cells are cultured on high stiffness hydrogels, which is exacerbated by loss of GORAB. These results indicate that vimentin IFs contribute to the structural stability of the Golgi apparatus, and suggest a role for GORAB in this process.

## Introduction

The Golgi apparatus has a characteristic structure in most eukaryotes, comprising a series of cisternae layered on top of each other to form the Golgi stack (Lowe, 2011; Shorter and Warren, 2002). In most vertebrate cells, the stacks are connected laterally to form the Golgi ribbon, which is usually positioned next to the centrosome by virtue of attachment to microtubules and the minus-end directed microtubule motor protein dynein (Brownhill et al., 2009; Mascanzoni et al., 2022; Thyberg and Moskalewski, 1999). The Golgi is a polarized structure, comprising an entry (cis-) face for cargo entering from the endoplasmic reticulum (Brandizzi and Barlowe, 2013), and an exit (trans-) face from where cargo is transported to various downstream destinations (Di Martino et al., 2019). Cargo transiting the Golgi is exposed to a series of resident cisternal enzymes that sequentially modify the cargo, most notably at the level of glycosylation, as it moves from one side to the other (Glick and Nakano, 2009; Stanley, 2011; Welch and Munro, 2019). In addition to microtubules, the Golgi is also intimately associated with the actin cytoskeleton, which in vertebrate cells is not important for ribbon positioning (Egea et al., 2013). Instead, it functions at a more local level, contributing to formation of transport vesicles and linking of stacks into the Golgi ribbon (Chakrabarti et al., 2021). Actin contributes to the mechanical rigidity of Golgi membranes, which is important for the membrane deformation events that occur during transport vesicle formation (Guet et al., 2014). Actin can also couple to microtubules through the formin protein FHDC1/IFN1, which is thought to coordinate actin and microtubule dynamics during Golgi ribbon formation (Copeland et al., 2016).

In contrast to microtubules and actin, much less is known about the association of the Golgi apparatus with the third major class of cytoskeletal element, intermediate filaments (IFs). IFs are present in all metazoans and are comprised of different proteins, depending on cell type and subcellular location (Etienne-Manneville, 2018; Herrmann et al., 2009). They provide important structural integrity to cells and contribute to a number of dynamic cell processes including cell migration, cell division and apoptosis (Etienne-Manneville, 2018; Herrmann et al., 2009). IFs are important for providing mechanical integrity to a number of cellular organelles, most strikingly in the case of the nucleus, where nuclear lamins play a key role (Gruenbaum and Foisner, 2015). Cytoplasmic vimentin IFs also contribute to the nuclear integrity by forming a cage around the nucleus that is mechanically protective (Patteson et al., 2019). A recent study has revealed that vimentin IFs can also associate with the endoplasmic reticulum and help maintain its morphological organization, as well as concentrate endolysosomes in the perinuclear area (Cremer et al., 2022). Vimentin cages can form around aggresomes to control protein degradation (Morrow and Moore, 2020; Morrow et al., 2020), lipid droplets (Franke et al., 1987), melanophores (Chang et al., 2009) and vimentin IFs also contribute to mitochondrial position and shape (Schwarz and Leube, 2016). In terms of the Golgi apparatus, previous work has shown that vimentin IFs can bind to Golgi membranes through the *cis*-Golgi localised peripheral protein forminotransferase cyclodeaminase (FTCD, also known as 58K) (Gao and Sztul, 2001; Gao et al., 2002). However, the functional significance of this association remains unclear. It is also unknown whether the Golgi can physically associate with IFs in other ways, and any functional relationship between IFs and the Golgi apparatus remains poorly defined.

The coiled-coil Golgi protein GORAB, which is mutated in the skin and bone disorder gerodermia osteodysplastica (GO) (Hennies et al., 2008), forms discrete regions or domains at the *trans*-Golgi network (TGN) (Witkos et al., 2019). GORAB associates through SCYL1 and ARF1 with the COPI vesicle coat complex and has been proposed to function as a scaffold for COPI vesicle formation at the TGN (Witkos et al., 2019). GORAB is also present at the centrioles, where it binds the cartwheel protein SAS-6 (Kovacs et al., 2018). GORAB binds SAS-6 as a monomer, whereas at the TGN it is present as a dimer (Fatalska et al., 2021). The function of GORAB at the centriole is unclear, but it may act as a physical scaffold, analogous to its proposed function at the TGN. In this study we report that GORAB can physically associate with vimentin IFs, and examine the relationship between vimentin IFs and Golgi architecture. Our results indicate that vimentin IFs are important to stabilize Golgi structure in mammalian cells, which is evident upon chemical perturbation of Golgi dynamics or culture in high mechanical stiffness. In contrast, GORAB does not play a major role in Golgi integrity, although it appears to sensitize the Golgi to the loss of vimentin IFs. These observations reveal a previously unappreciated role for vimentin IFs in providing structural stability to the Golgi apparatus.

## Results

### GORAB can physically associate with vimentin intermediate filaments

We have previously shown that GORAB is present at the TGN in discrete membrane domains and that it can scaffold COPI assembly for retrograde vesicle transport (Witkos et al., 2019). However, the mechanisms underlying GORAB function remain poorly defined, and it is also unclear whether GORAB has additional functions at the TGN. To gain more insight into possible GORAB functions, we performed proximity biotinylation using GORAB fused to the biotin ligase BirA to identify closely associated proteins (Roux et al., 2012). Stable expression of GORAB-BirA in HeLaM cells and human dermal fibroblasts, where it correctly localized to the Golgi (Fig **S1A**,**D**), resulted in biotinylation of Golgi-associated proteins, as revealed by immunofluorescence microscopy (Fig **S1B,E**) and Western blot (Fig **S1C,F**). Mass spectrometry identified a number of biotinylated proteins, including GORAB itself, the COPI subunit γ2-COP and Scyl1 (**Supplementary Tables 1 and 2**), consistent with our earlier work on GORAB and COPI at the TGN and a previous study on Drosophila GORAB (Kovacs et al., 2018; Witkos et al., 2019). A number of other Golgi-associated proteins were also identified, together with various cytoskeletal proteins and proteins previously identified in other Bio-ID screens, which are likely non-specific (Roux et al., 2012). Vimentin, although often found as a contaminant in proteomics experiments (www.crapome.org), was one the most abundant biotinylated proteins in both cell types, prompting further investigation.

To explore a possible physical association of GORAB with vimentin, GORAB was stably over-expressed in RPE-1 cells and colocalization with vimentin performed. As shown in Fig **S2A,B**, when expressed at high levels, GORAB was able to form extensive filaments extending throughout the cytoplasm, which co-aligned with vimentin IFs. A similar observation was reported previously upon GORAB over-expression in HeLa cells and fibroblasts (Egerer et al., 2015). The GORAB filaments did not colocalize with microtubules or actin filaments (Fig **S2B**). The co-alignment of GORAB with vimentin IFs was preserved following nocodazole treatment (Fig **S2B**), which retracts the IF network as reported previously (Yoon et al., 1998), as well as after collapse of IFs by expression of a vim_1-138_ dominant-negative mutant that impairs IF assembly (Fig **S2C**). GORAB fails to form filaments in SW13^−/−^ cells that lack all cytoplasmic IFs (Hedberg and Chen, 1986), instead forming ‘squiggles’, short filamentous elements reminiscent of structures seen during cytoplasmic IF assembly (Fig **S2D**). Thus, the formation of extended GORAB filaments is dependent upon the presence of pre-existing vimentin IFs. The formation of GORAB filaments also requires GORAB oligomerization, as the previously described K190del mutant, which cannot self-associate (Witkos et al., 2019), is deficient in oligomerization, even in the presence of an intact vimentin IF network (Fig **S2E**). These results suggest that vimentin IFs can scaffold assembly of GORAB filaments, consistent with a physical interaction between the proteins, at least under the conditions of these experiments. Imaging of SW13^−/−^ cells with residual vimentin expression supports this conclusion, with GORAB apparently incorporated into or closely aligned with the fragmented IFs seen in these cells (Fig **S2F**). It is worth noting that over-expression or loss of GORAB did not alter the vimentin IF network in wild-type cells, suggesting that GORAB does not play a role in organization of IFs (Fig **S2A,B** and Fig **S3A,B**). Together, these results indicate that GORAB has the ability to physically associate with vimentin IFs.

### Physical proximity of the Golgi apparatus to vimentin intermediate filaments

To better understand the spatial relationship of GORAB, and the Golgi apparatus more generally, with vimentin IFs, confocal microscopy was performed. As expected, vimentin IFs were observed to form a network extending throughout the cytoplasm, with an apparent enrichment in the perinuclear region where the Golgi is usually localised (Fig **1**). Confocal imaging of mouse embryonic fibroblasts (MEFs) revealed that both GORAB and the *cis*-Golgi marker GM130 are not only in close physical proximity to the IF filaments, but apparently embedded in a meshwork of filaments, as observed in various tilted 3D projections (Fig **1A,B**). Further analysis was performed on human skin fibroblasts using confocal microscopy followed by deconvolution to enhance the signal/noise ratio and better discern the spatial relationship of the TGN with vimentin IFs. This revealed that the TGN is also closely associated with vimentin IFs (Fig **1C)**. The TGN occasionally can be seen to co-localise with IFs, but more frequently it is aligned with IFs, with the IFs juxtaposed on one or both sides of the TGN membrane (Fig **1C**). GORAB puncta can be seen along along the TGN, adjacent to the IFs. Similar results were obtained in RPE-1 and HeLa cells, where again the TGN membrane and GORAB were found co-localised with or juxtaposed on one or both sides by IFs (Fig **1D,E**). Because other cytoskeletal elements such as microtubules are also enriched in the perinuclear region of the cell, and IFs associate with both microtubules and actin filaments, these observations do not necessarily show that the Golgi is physically associated with vimentin IFs. Nevertheless, they reveal an intimate spatial relationship between them.

**Figure 1.**
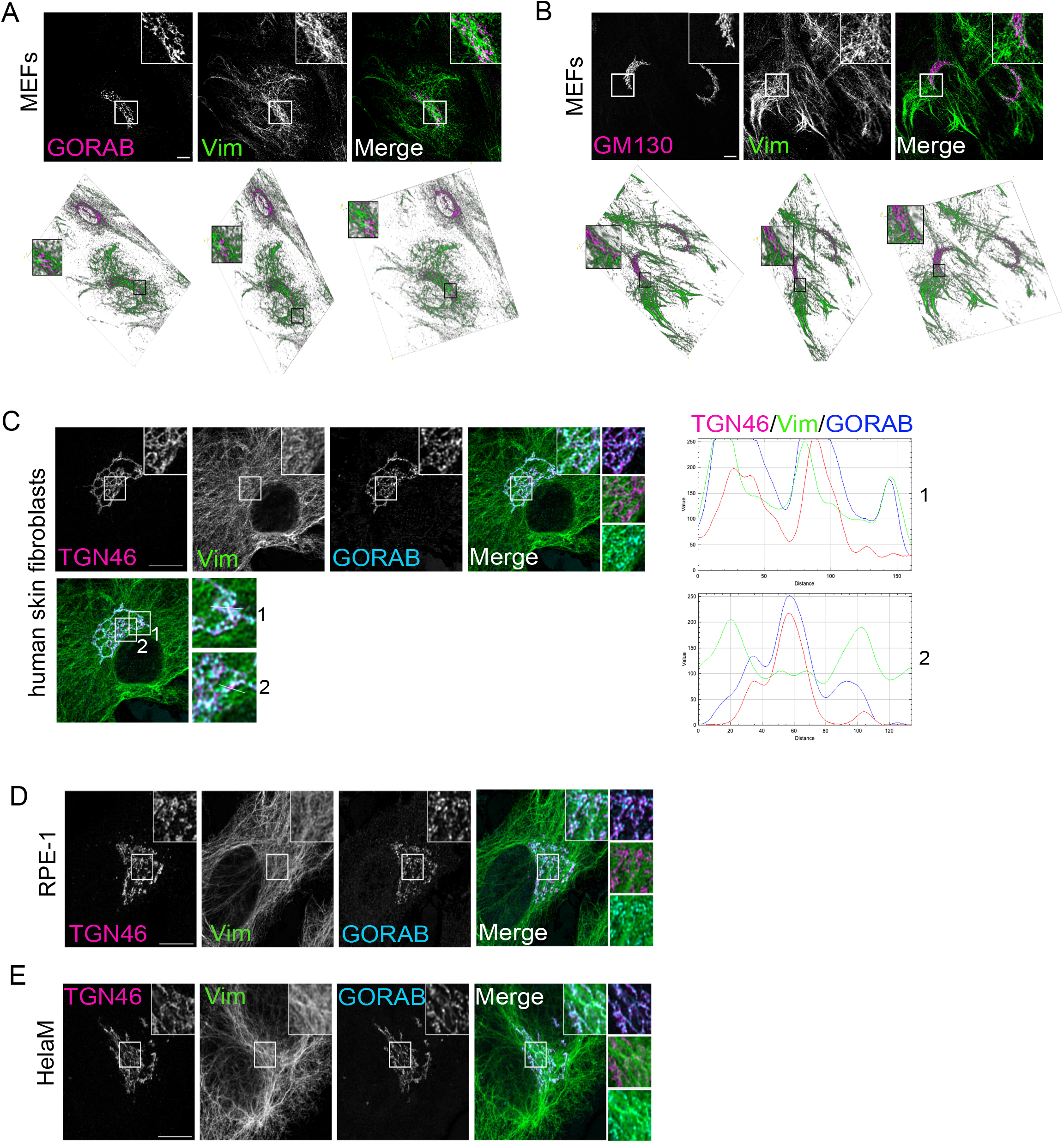
Spatial relationship of GORAB, GM130, TGN46 and vimentin filaments in mouse and human cells. **A,B)** 2D (z stack maximal intensity projection) and 3D confocal images of WT MEFs stained with antibodies to vimentin (Vim) and either GORAB (A) or GM130 (B). Scale bar, 10 μm; **C,D,E)** Deconvolved confocal z stack maximal intensity projection images of human skin fibroblasts (C), RPE-1 (D) or HelaM (E) cells stained with GORAB, TGN46 and vimentin antibodies. The white lines in C depict the areas taken for calculating RGB fluorescence intensity profile plots in human skin fibroblasts, shown on the right. Scale bars, 10 μm.

### Loss of GORAB or vimentin has a modest effect on Golgi architecture at steady state

We next wanted to test whether vimentin IFs play a role in Golgi integrity, positioning or morphology. In parallel, we investigated whether GORAB may also play a role in these processes, independently or in conjunction with vimentin. For this purpose, we used vimentin null MEFs that were obtained from a previously generated vimentin knockout mouse (Colucci-Guyon et al., 1994), and performed CRISPR/Cas9-mediated knockout of GORAB in these cells to make a double knockout, or in wild-type MEFs, to make a single knockout cell line. Western blotting and immunofluorescence microscopy confirmed the knockout of vimentin and GORAB in the single and double knockout lines, as expected (Fig **2A-C**). There was no major change in the morphology or positioning of the ER (labelled using PDI), the ER to Golgi intermediate compartment (ERGIC, labelled using Scyl1), or the Golgi apparatus (labelled using TGN marker syntaxin 6 (Stx6)) in the single and double knockout cells (Fig **2D**). Confocal imaging of the *cis*-Golgi marker GM130 indicated the Golgi ribbon was intact in all knockout lines (Fig **2E,F**). Transmission electron microscopy was next used to examine Golgi ultrastructure in the knockout cells. This revealed a slight reduction in cisternal number and a shortening of the cisternal length in all three knockout lines (Fig **3A,B**). Consistent with the presence of stacked cisternae in the knockout cells, *cis-trans* polarity of the Golgi, assessed by immunofluorescence microscopy of nocodazole-induced Golgi ministacks, was maintained (Fig **3C,D**). Golgi polarity was also maintained in the SW13−/− cells lacking intermediate filaments (Fig **S3C**). Thus, although there are changes in Golgi morphology in the knockout cells, these are fairly modest, and ribbon integrity appears unaffected. Together, these results indicate that GORAB and vimentin do not play a major role in Golgi organization, polarity or positioning under steady state conditions.

**Figure 2.**
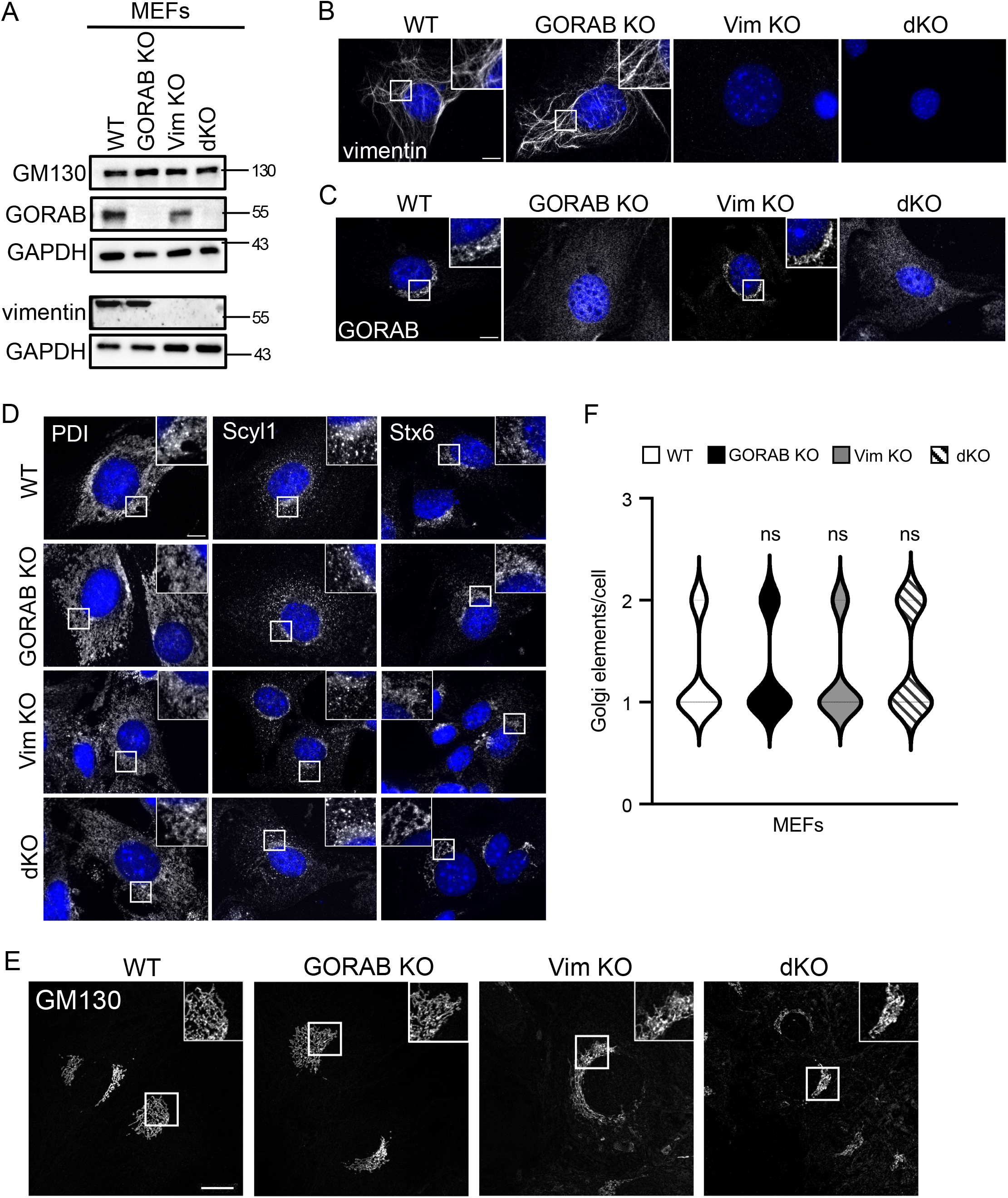
Generation of GORAB KO in WT and Vim KO MEFs and effects upon ER and Golgi morphology. **A)** Western blot analysis of GORAB, vimentin and GM130 expression levels in WT, GORAB KO, Vim KO and dKO MEFs; **B, C)** Immunofluorescence of vimentin (B) in WT and Vim KO MEFs and of GORAB in WT, GORAB KO, Vim KO and dKO MEFs. Scale bar, 10 μm; **D)** Immunofluorescence of ER (PDI), ERGIC (Scyl1) and TGN (Stx6) in WT, GORAB KO, Vim KO and dKO MEFs. Scale bar, 10 μm; **E)** Confocal images (z stack maximal intensity projection) in WT, GORAB KO, Vim KO and dKO MEFs. Scale bar, 10 μm; **F)** Analysis of the numbers of *cis*-Golgi (GM130) elements in WT, GORAB KO, Vim KO and dKO MEFs. In each case comparisons are shown for the specified KO cell line versus WT cells, with statistical significance calculated using a non-parametric one-way ANOVA (Kruskal-Wallis test).

**Figure 3.**
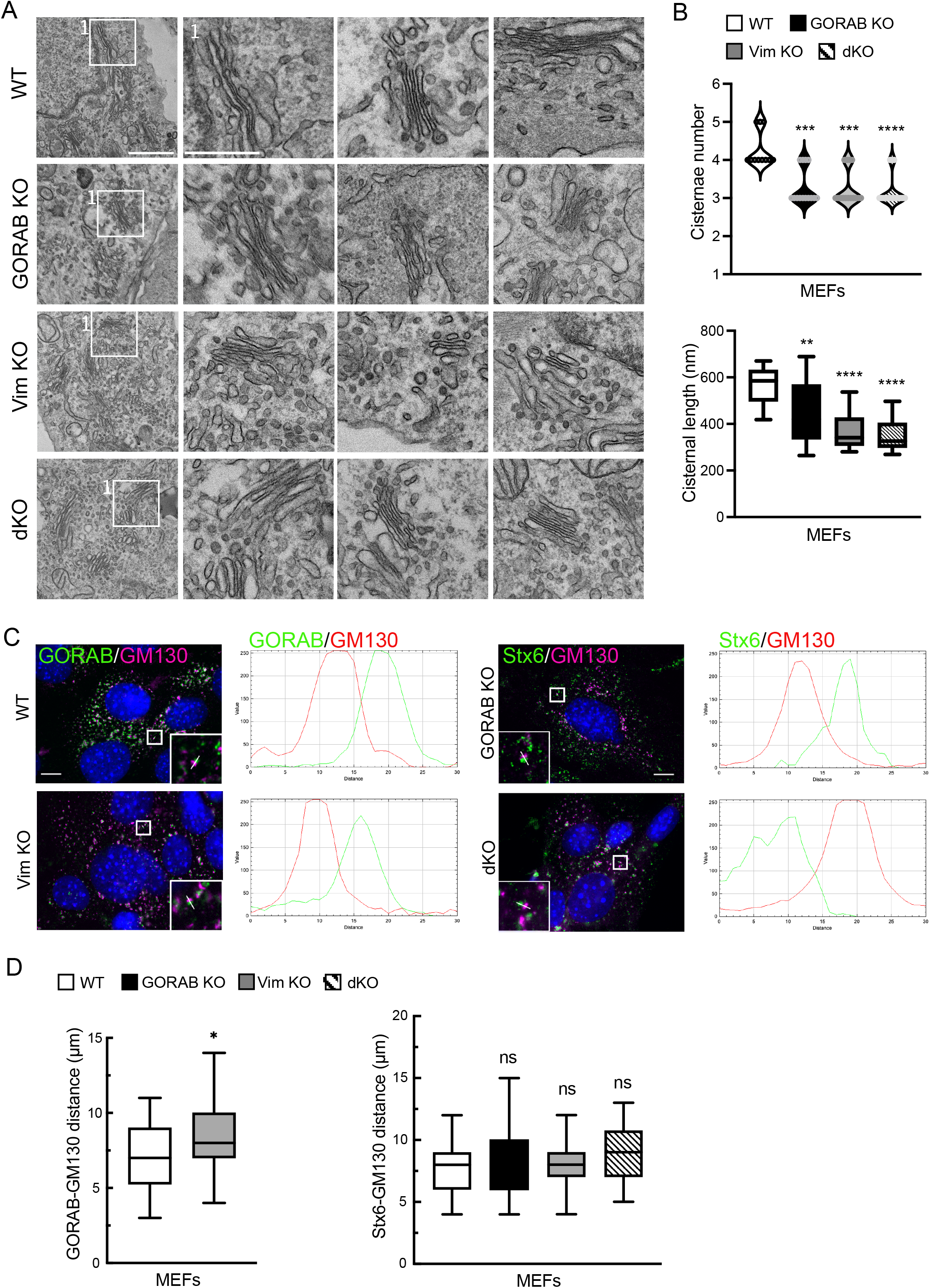
Ultrastructure and polarisation of the Golgi in WT, GORAB KO, Vim KO and dKO MEFs. **A)** TEM images of the Golgi in WT, GORAB KO, Vim KO and dKO MEFs. Scale bar, 500 nm (left column) or 250 nm (right columns); **B)** Analysis of cisternae number and length from TEM images in WT, GORAB KO, Vim KO and dKO MEFs. In each case comparisons are shown for the specified KO cell line versus WT cells, with statistical significance calculated using a non-parametric one-way ANOVA (Kruskal-Wallis test) (top) or one-way ANOVA with post-hoc Dunnett test (bottom); **C)** Immunofluorescence of *cis*-Golgi (GM130) and TGN (GORAB or Stx6) polarization in WT, GORAB KO, Vim KO and dKO MEFs 2 h after nocodazole treatment. White line depicts the area taken for calculating RGB fluorescence intensity profile plots. Scale bar, 10 μm; **D)** Analysis of *cis*- and TGN Golgi stack distance in WT, GORAB KO, Vim KO and dKO MEFs. In each case comparisons are shown for the specified KO cell line versus WT cells, with statistical significance calculated using an unpaired t-test (left) or one-way ANOVA with post-hoc Dunnett test.

### Loss of vimentin causes faster disassembly and slower reassembly of the Golgi apparatus upon chemical perturbation

To further explore whether vimentin IFs play a structural role at the Golgi, we used the drug brefeldin A (BFA) to trigger Golgi disassembly in control or knockout MEFs. BFA causes disassembly of the Golgi apparatus by inhibiting ARF guanine nucleotide exchange activity, which can be reversed by washing out the drug (Klausner et al., 1992). We reasoned that if vimentin IFs (or GORAB) provide underlying mechanical support to the Golgi, then their loss would result in a faster Golgi fragmentation upon BFA treatment. For these experiments we focused on the initial stages of Golgi disassembly, occurring over 10 minutes. As shown in Fig **4A,C** and Fig **S4A,B**, as expected, treatment of wild-type MEFs caused disassembly of the Golgi, visualized using antibodies to GORAB, and markers to the *cis*- (Golgin-84, GM130) and *trans*- (Stx6) Golgi. Disassembly was quantified by measuring immunofluorescence signal above threshold, which confirmed the progressive loss of Golgi signal over the first 10 minutes of incubation (Fig **4B,D**). Loss of vimentin accelerated disassembly of the Golgi apparatus, as indicated by all Golgi markers (Fig **4A-D**). A similar result was seen with the double knockout cells, whereas loss of GORAB alone did not affect the rate of disassembly (Fig **4D** and Fig **S4A,B**). To further assess the role of intermediate filaments in Golgi stability, the experiment was repeated in SW13^−/−^ cells lacking all cytoplasmic IFs. As seen in the MEFs, there was a more rapid fragmentation of the Golgi apparatus upon BFA treatment in the SW13^−/−^ cells compared to wild-type SW13^+/+^ cells (Fig **S5A,B**). The microtubule network appeared normal in the vimentin knockout MEFs (Fig **4E**), and this was also the case in SW13^−/−^ cells (Fig **S5C**), indicating that the changes in Golgi stability and GORAB localization are not due to indirect effects on the microtubule cytoskeleton. Similarly, we did not see any changes in filamentous actin organization in the vimentin or double knockout MEFs or SW13^−/−^ cells, or in focal adhesions, again arguing against indirect effects via the actin cytoskeleton when vimentin IFs are missing (Fig **S6A,B**).

**Figure 4.**
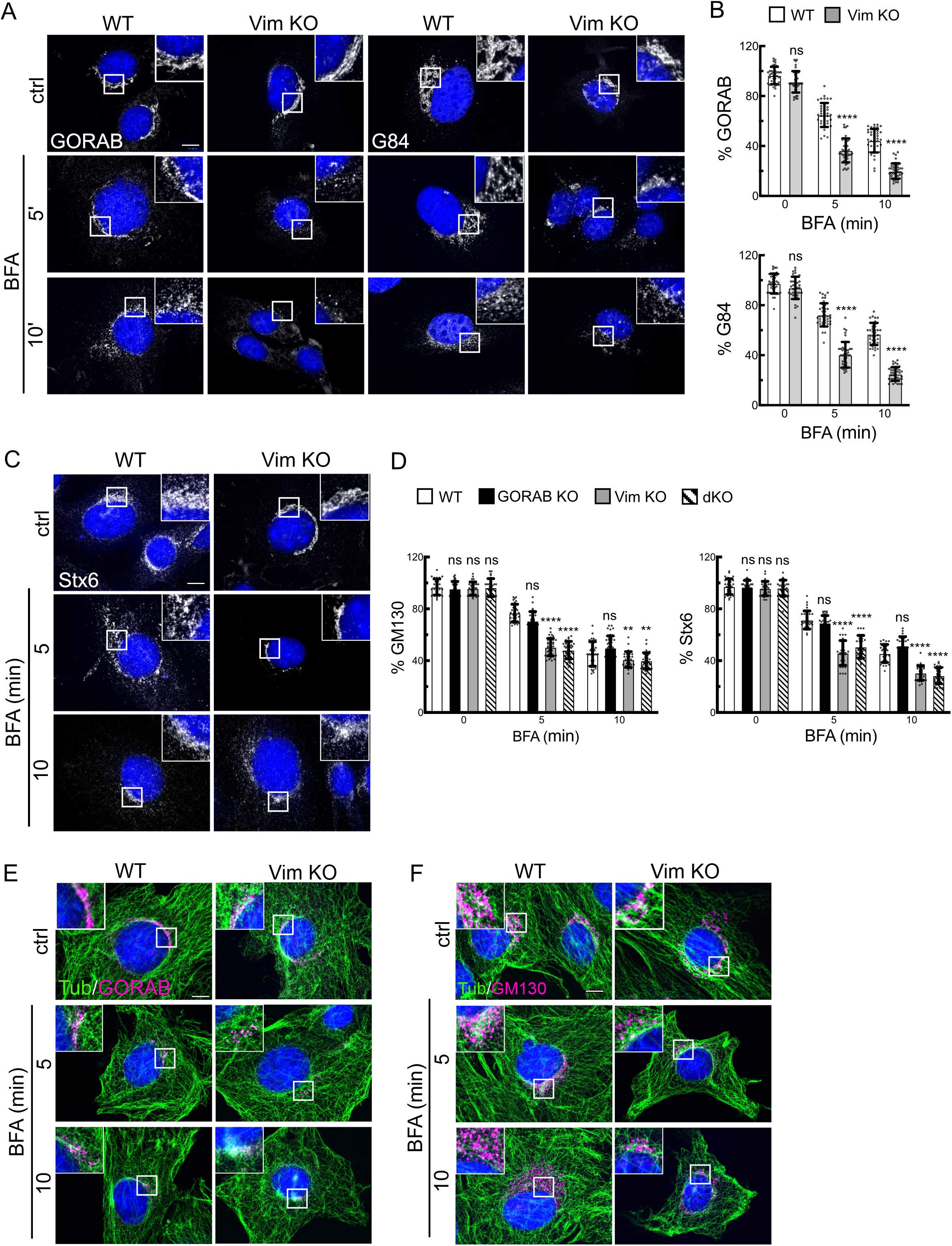
Golgi fragmentation in WT and Vim KO MEFs treated with BFA. Cells were incubated in the presence of BFA (5 μg/ml) at 37°C for the indicated times prior to fixation and immunostaining. **A,C)** Immunofluorescence of GORAB -Golgin-84 (A) and GORAB– Syntaxin6 (C) in WT and Vim KO MEFs; **B,D)** Quantification of GORAB and G84 (B) and GM130 and Stx6 (F) fluorescence intensities in WT, GORAB KO, Vim KO and dKO MEFs treated for 5 and 10 min with BFA. At each timepoint, comparisons are shown for the specified KO cell line versus WT cells. Statistical significance between WT and KO cells was calculated using an unpaired t-test (B) or two-way ANOVA with post-hoc Dunnett test (D); **E)** IF staining of GORAB with tubulin in MEFs WT and VimKO; **F)** Immunofluorescence of GM130 with tubulin in WT and Vim KO MEFs. Scale bar, 10 μm.

To further assess a possible structural role for vimentin IFs in Golgi organization, the rate of Golgi reassembly following BFA washout was analyzed. The MEFs were treated with BFA for 90 minutes to induce complete Golgi fragmentation, which was seen in wild-type and all three knockout cell lines (Fig **5A,B,D**, 0 min washout). GM130 was present in puncta corresponding to ERES as reported previously (Mardones et al., 2006), while Stx6 was concentrated at the centrosome, in line with previous studies showing TGN concentration at the centrosome upon BFA treatment of rodent cells (Reaves and Banting, 1992). After 30 minutes washout of BFA, Golgi elements containing the markers GM130 and Stx6 started to concentrate in the perinuclear region of wild-type cells, consistent with the reassembly of the Golgi ribbon, which was further progressed at 45 min washout (Fig **5C,D**). Interestingly, the appearance of GORAB in the reforming Golgi ribbon appeared slower than the other markers (Fig **5A**). Reassembly in the vimentin knockout cells was significantly delayed compared to wild-type MEFs, as indicated by GORAB (Fig **5A,B**), GM130 (Fig **5C,E**) and Stx6 (Fig **5D,F**). A similar result was obtained with the double knockout cells, whereas loss of GORAB alone did not affect the kinetics of Golgi reassembly (Fig **5E,F** and Fig **S4C**). A similar delay in Golgi reassembly was also seen in SW13^−/−^ cells compared to wild-type SW13^+/+^ cells, confirming the effect is due to loss of IFs (Fig **S5E,F**). The Golgi ultimately fully reformed at later timepoints in all cell lines, indicating that the loss of vimentin only slows, as opposed to arrests, Golgi reassembly under these conditions (Fig **S4D**). Together, the results support a role for vimentin in scaffolding Golgi membranes such that they undergo faster BFA-induced disassembly and slower reassembly in its absence.

**Figure 5.**
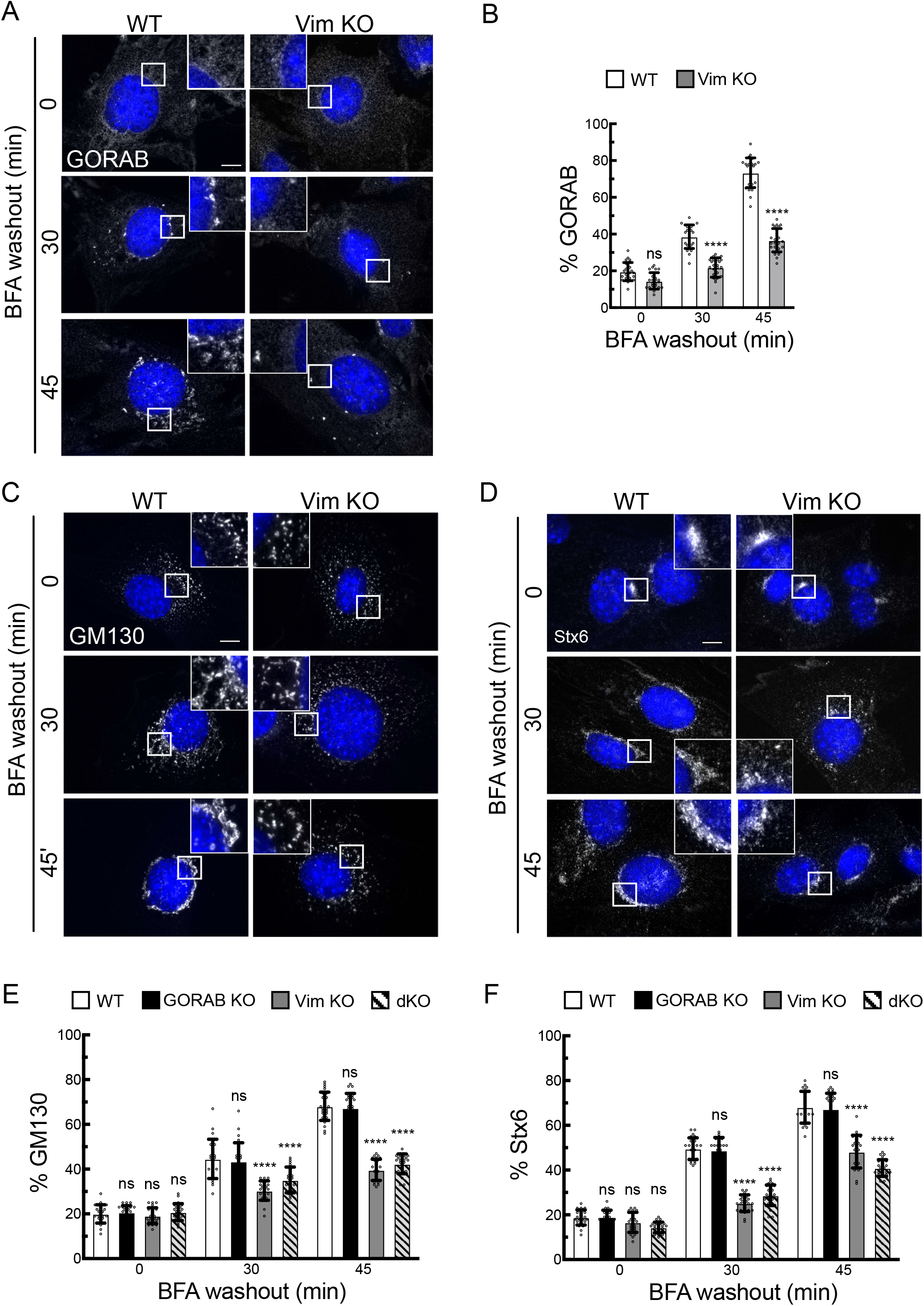
Golgi reassembly in WT and Vim KO MEFs upon BFA washout. Cells were incubated with BFA (5 μg/ml) for 90 min at 37°C, washed and incubated in fresh medium for the indicated time points, prior to fixation and immunostaining. **A)** Immunofluorescence of GORAB in WT and Vim KO MEFs after BFA washout. Scale bar, 10 μm; **B)** Quantification of GORAB reassembly in WT and Vim KO MEFs. Comparison between groups was performed with an unpaired t-test; **C,D)** Immunofluorescence of GM130 (C) and Stx6 (D) in WT and Vim KO MEFs after BFA washout. Scale bar, 10 μm; **E-F)** Quantification of GM130 (E) and Stx6 (D) reassembly in WT, GORAB KO, Vim KO and dKO MEFs. At each timepoint comparisons are shown for the specified KO cell line versus WT cells, with statistical significance calculated using a two-way ANOVA with post-hoc Dunnett test.

### Loss of vimentin destabilizes Golgi structure in a high stiffness environment

The vimentin IF network provides mechanical support to cells, but the extent to which can mechanically support individual organelles is less clear. To examine whether vimentin IFs may play a role in providing mechanical support to the Golgi apparatus, cells were cultured on hydrogel substrates of differing stiffness, ranging from 2 kPa (soft) to 170 kPa (stiff). In terms of physiological comparison, these would of the order seen in soft tissues such as brain and adipose versus stiffer tissues such as precalcified bone and cartilage, respectively (Discher et al., 2009; Guimarães et al., 2020). In wild-type MEFs, the Golgi appeared unaffected in terms of ribbon morphology on the different substrates (Fig **6A,C**). This indicates that, at least in MEFs, the Golgi does not undergo significant changes in its organization when substrate stiffness is altered. A similar result was seen with GORAB knockout MEFs (Fig **6B,C**). However, in vimentin knockout MEFs, there was a progressive increase in fragmentation of the Golgi ribbon as substrate stiffness increased (Fig **6A,C**). The extent of fragmentation was greater in the double knockout cells compared to vimentin only knockout, an effect that was particularly evident at lower stiffnesses (Fig **6B,C**). These observations suggest that vimentin IFs provide mechanical support to the Golgi ribbon, which is important in higher stiffness environments, and that GORAB also contribute to the mechanical integrity of the Golgi ribbon.

**Figure 6.**
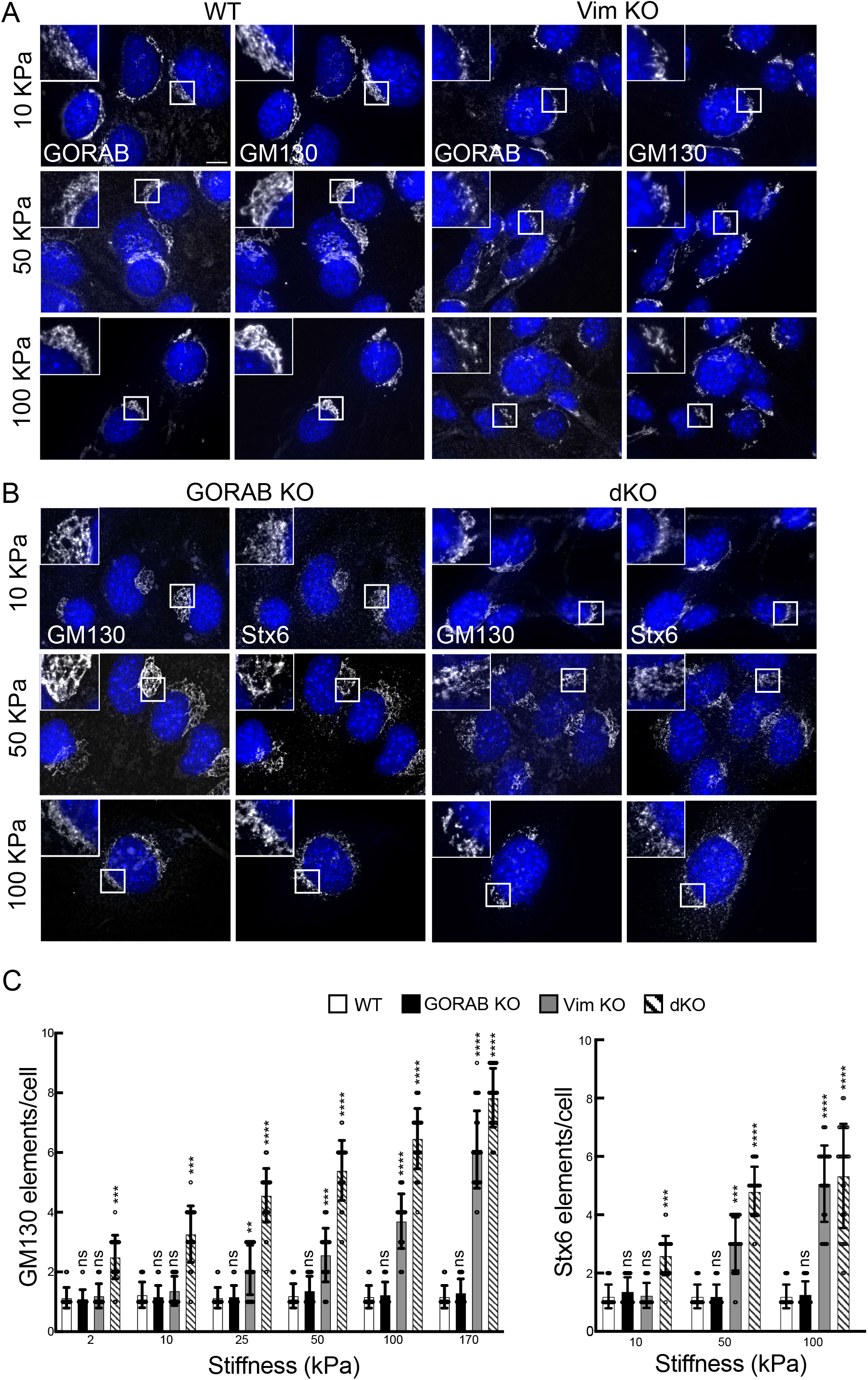
Golgi morphology in the KO MEFs in response to different stiffness substrates. The indicated cells were plated on elastic matrigel coated coverslips for 48 h prior to fixation and immunostaining. **A)** Cells were labelled with antibodies to GORAB and GM130; **B)** GM130 and Stx6. Scale bar, 10 μm; **C)** Analysis of the numbers of *cis*-Golgi (GM130) and TGN (Stx6) elements in WT, GORAB KO, Vim KO and dKO MEFs upon culture in different stiffness. At each stiffness comparisons are shown for the specified KO cell line versus WT cells, with statistical significance calculated using a non-parametric one-way ANOVA (Kruskal-Wallis test).

### Discussion

Although cytoplasmic IFs are known to provide mechanical support to cells, their functional relationship with organelles is less well understood. In particular, the extent to which IFs associate with the Golgi apparatus, and the role of any association, remains poorly defined. Our data indicate a close spatial relationship between the Golgi and vimentin IFs, and that loss of vimentin reduces the structural integrity of the Golgi ribbon. How vimentin IFs provide support to the Golgi remain to be determined, but an attractive hypothesis is that they form a cage or meshwork around the organelle, which provides a mechano-protective function. This would be analogous to what has been reported for the nucleus and other cytoplasmic organelles (Cremer et al., 2022; Etienne-Manneville, 2018). The Golgi ribbon, by virtue of its localization adjacent to the nucleus, may be embedded within the IF meshwork that forms a cage around the nucleus (Cremer et al., 2022; Etienne-Manneville, 2018). This would certainly be consistent with our imaging data. The integrity of the Golgi may therefore be controlled by the same network of vimentin IFs that maintain nuclear integrity and that of other perinuclear organelles, allowing for a coordinated control of organelle mechanics.

Whether the Golgi binds directly to vimentin IFs, and the possible role of any physical interaction, remains poorly defined. Previous studies identified the Golgi-localised metabolic enzyme FTCD as a vimentin binding protein (Gao and Sztul, 2001), but the significance of this interaction for Golgi integrity or function remains unclear. FTCD over-expression alters vimentin IF organization as opposed to affecting Golgi morphology per se, and it was proposed that the Golgi may act as a reservoir for FTCD that in turn may modulate vimentin dynamics (Gao and Sztul, 2001; Gao et al., 2002). In this study, we identified TGN-localized coiled-coil protein GORAB as a possible vimentin interactor. As previously reported for FTCD, over-expressed GORAB associates with vimentin IFs, although in contrast to FTCD, it does not appear to affect the organization of IFs. Loss of GORAB also did not affect Golgi ribbon morphology, although in high stiffness it did sensitize the Golgi to loss of vimentin. Whether GORAB can bind vimentin at endogenous levels, and whether this interaction can provide a physical link between the TGN membrane and vimentin IFs, remains an open question. It is interesting to note that GORAB oligomerization is necessary for binding to vimentin IFs, and it is also required for the formation of TGN-associated GORAB domains involved in COPI vesicle traffic (Witkos et al., 2019). Hence, GORAB oligomerization may control the various functions of the protein, as proposed by Fatalaska et al, who found that centriolar GORAB is present as a monomer, whereas it is a dimer at the TGN (Fatalska et al., 2021). Interestingly, the K190 deletion, a causative mutation in GO that disrupts GORAB olgomerisation, abolishes both TGN domain formation and IF binding, raising the possibility that IF binding could also be disrupted in GO. FTCD was previously shown to bind SCYL1 (Burman et al., 2010), which is a binding partner of GORAB (Di et al., 2003; Witkos et al., 2019), raising the additional possibility that GORAB may form a common complex with FTCD that can bind to vimentin IFs. It is also of course possible, that the Golgi may bind to vimentin via other proteins, which remain to be identified.

Previous studies have shown that the Golgi apparatus can act in a mechanoresponsive manner, with changes in cell mechanics resulting in altered Golgi function (Guet et al., 2014; Romani et al., 2019). Using a micro-rheology approach Guet et al showed that Golgi membrane tension can influence the generation of Golgi-derived Rab6-positive vesicles (Guet et al., 2014). They could also show that Golgi membrane tension is dependent upon the actin cytoskeleton. Similarly, it was recently shown that ER-Golgi traffic of the SREBP transcription factor, which controls lipid metabolism, is also mechanoresponsive, again downstream of the actin cytoskeleton (Romani et al., 2019). The actin cytoskeleton can therefore dictate the functional response of the Golgi apparatus to changes in cell mechanics. Our data suggest that vimentin IFs may also play a role in this response. For example, we may hypothesize that by providing mechanical support to Golgi membranes, vimentin IFs could influence rates of vesicle formation at this organelle, a process which is dependent upon the extent of physical deformation of the Golgi membrane. Similarly, maintaining the integrity of the Golgi ribbon is likely to have important functional consequences considering an intact Golgi ribbon is required to maintain optimum protein glycosylation (Puthenveedu et al., 2006; Xiang et al., 2013), guide polarized secretion (Ravichandran et al., 2020; Yadav et al., 2009), and can influence the progression through the cell cycle (Mascanzoni et al., 2022; Rabouille and Kondylis, 2007; Sutterlin et al., 2002). It will be interesting to investigate the role of vimentin IFs in maintaining these and other aspects of Golgi function, as well as the extent of functional interplay between IFs and the actin cytoskeleton at the Golgi.

An apparent conundrum from our experiments is that loss of vimentin strongly affected the integrity of the Golgi ribbon in cells grown on stiff hydrogels, but appeared dispensable for cells growing at steady-state on plastic or glass, which are orders of magnitude stiffer than the highest stiffness gel we used (~1 GPa versus 170 kPa). These results would suggest the nature of the substrate, as well as its inherent stiffness, are important for the responsiveness of the Golgi membranes to loss of vimentin. Glass and plastic lie far beyond the physiological range of stiffness found within tissues, and they are also clearly chemically distinct to the extracellular matrix found within tissues and also the hydrogels we used. Considering vimentin IFs bind to focal adhesions and there is functional cross-talk between these structures (Kreis et al., 2005; Leube et al., 2015), it is perhaps not surprising that changes in dependencies upon vimentin will differ depending on the chemical nature of the cellular environment. In addition, it is known that changes in material chemistry can influence the conformation of fibronectin displayed to cells (Garcia et al., 1999; Rico et al., 2009), which could also contribute to the changes we observe between glass (with passively adsorbed fibronectin from the medium), and hydrogels (with covalently attached collagen). By using the same chemical substrate in our hydrogels, we were able to differentiate between effects due to cell substrate versus stiffness, and our results indicate a clear dependency upon vimentin for Golgi stability at high but not low stiffness. Clearly further studies will be interesting to perform using both 2D and 3D culture on substrates of different composition, assessing the role of vimentin in maintaining Golgi integrity under the different conditions.

## Materials and methods

### Antibodies

The following antibodies were used in this study: rabbit anti-GORAB (Proteintech 17798-1-AP), 1:500 (WB) and 1:100 (IF); rat anti-vimentin (R&D MAB21052), 1:500 (WB) and 1:100 dilution (IF); for WB and IF; mouse anti-vimentin (Santa Cruz V9), 1:100 (IF); mouse anti-vinculin (Sigma V9131), 1:200 (IF); rabbit anti-GM130 (MLO7 anti-N73pep, (Nakamura et al., 1997)), 1:1000 (WB) and 1:100 (IF); mouse anti-GM130 (BD Bioscience 610822), 1:1000 (WB) and 1:100 (IF); mouse anti-Stx6 (from Andrew Peden, University of Sheffield), 1:50 (IF); sheep anti-TGN46 (from Vas Ponnambalam, University of Leeds), 1:300 (IF); mouse anti-PDI (Enzo ADI-SPA-891), 1:1000 (WB) and 1:100 (IF); mouse anti-⍰-tubulin, (from Keith Gull, University of Oxford), 1:1000 (WB) and 1:100 (IF); mouse anti-GAPDH (SantaCruz sc-365062), 1:2000 (WB); mouse anti-Myc (Cell Signalling, 9B11), 1:1000 (IF). Anti-Scyl1 antibodies were raised in sheep against recombinant GST-tagged full-length human Scyl1 and affinity purified against the recombinant protein. Sheep anti-Scyl1 was used at 1:500 (WB) and 1:50 (IF). Actin was labelled with Alexa 594-conjugated phalloidin (ThermoFisher), 1:1000. HRP-conjugated secondary antibodies for immunoblotting were from Sigma (A4416 and A0545, both at 1:1000). Fluorophore-conjugated secondary antibodies for immunofluorescence were from Jackson ImmunoReserach laboratories. HRP- and Alexa 488-conjugated streptavidin for BioID experiments were from ThermoFisher.

### Cell Culture

HeLaM, hTERT-RPE-1, and wild-type MEF cells were from American Type Culture Collection (ATCC, Manassas, VA). Vimentin KO MEFs from a vimentin null mouse model were a kind gift from Dr. Sara Koester (University of Gottingen) (Colucci-Guyon et al., 1994). Human skin fibroblasts derived from healthy individuals and from GO patients were obtained from Dr May Tassabehji (University of Manchester, UK) and from the Cell Line and DNA Bank from Patients affected by Genetic Diseases (Genova, Italy). SW13 clone 1 (vim^+/+^) and clone 2 (vim^−/−^) were kind gifts from Victoria Allan (University of Manchester). All cells apart from hTERT-RPE-1 were cultured in Glutamax DMEM (ThermoFisher 31966021) supplemented with 10% Fetal Bovine Serum (FBS), 50 U/ml penicillin and 50 μg/ml streptomycin. hTERT-RPE-1 cells were cultured in Ham’s F12 and DMEM 1:1 (ThermoFisher 11320033) with the same supplements. Cells were cultured at 37°C in 5% CO2.

### Growth of cells on polyacrylamide hydrogels

Polyacrylamide hydrogels with different substrate stiffnesses were prepared according to the established sandwich protocol described by (Tse and Engler, 2010). Glass slides were treated with dichlorodimethylsilane for 5 min to create a hydrophobic surface that prevents polyacrylamide attachment. Next, coverslips of 15 mm diameter were treated with 3-aminopropyltriethoxysilane and glutaraldehyde. The resulting amino-silanated coverslips were air-dried. Acrylamide (40%) and bis-acrylamide (2%) were mixed together in various relative concentrations to tune the resulting elasticities. Solutions of acrylamide/bis-acrylamide were crosslinked via a free radical polymerization reaction using ammonium persulfate and tetramethyldiethylenediamine. Immediately after the addition of crosslinkers, 20 μL of acrylamide/bis-acrylamide solution was sandwiched between the glass slide and the amino-silanated coverslip. All polyacrylamide gels were allowed to completely crosslink at room temperature for 2 h. Polyacrylamide gels were then washed twice in HEPES buffer (50 mM, pH 8.5) and sterilized with UV inside a biological safety cabinet. To allow adhesion, polyacrylamide gels were covalently crosslinked with human fibronectin. A solution of 0.2 μg/μL sufosuccinimidyl-6-(4 -azido-2 -nitrophenylamino)-hexanoate (sulfo-SANPAH) was freshly prepared in HEPES buffer with 1% dimethylsulfoxide. The sulfo-SANPAH solution was placed on gels and activated using 1 Joule of 254 nm UV light (Stratagene UV crosslinker). Afterwards, polyacrylamide gels were washed once with HEPES buffer and incubated with 10 μg/mL fibronectin in HEPES buffer overnight at 4°C. Polyacrylamide gels coated with fibronectin were used within one week of preparation to ensure mechanical integrity.

### Generation of cell lines stably expressing GORAB

All cDNA sequences were cloned into pXLG3 MCS (a modified version of pXLG3 with an expanded multiple cloning site) for lentivirus production. GORAB was tagged at the amino terminus with BirA*, mApple or GFP by cloning into the SpeI and XhoI or BamHI.sites of pXLG3 MCS. Lentiviruses were prepared as follows, based on a previously described method (Salmon and Trono, 2007). 24 h after seeding, HEK293 LTV (2.5 × 10^6^ cells per 10 cm dish) cells were co-transfected with 6 μg of the appropriate pXLG3 construct and 4.5 g psPAX2 and 3 μg pM2G vectors (packaging and envelope vectors respectively) using 27 μL of polyethylenemine mix. 24 h after transfection 100 μL of 1 M sodium butyrate was added. After 8 h the medium was changed to fresh DMEM supplemented with 10% HyClone FBS and 1 mM L-glutamine. 72 h after initial transfection, medium containing lentivirus was gathered, centrifuged at 4,000 rpm for 10 min at RT and the supernatant was filtered through a 0.45 μm filter (EMD Millipore, Billerica, MA). The lentivirus-containing medium was snap frozen in 1 mL aliquots and stored at −80°C. For lentiviral transduction, cells were plated in 10 cm dish containing 1 × 10^6^ cells and incubated with 1-3 mL of filtered medium containing lentivirus for 3 days in antibiotic-free medium. The efficiency of transduction was validated by immunofluorescence microscopy. For stable transduction, cells were trypsinised and seeded into a new 10 cm dish at 1:200,000 dilution and cells were grown in fully supplemented DMEM for a period of 10 to 14 days until distinct cell colonies were formed. The cell medium was removed, cells were washed with DMEM and pre-warmed trypsin and incubated for 3 min at 37°C. Colonies were picked using a sterile tip and propagated. The expression levels of tagged GORAB proteins were validated by immunofluorescence microscopy and western blotting.

### Generation of GORAB knockout MEFs using CRISPR-Cas9

GORAB was targeted using the abm custom All-In-One lentivirus system. Lentiviruses containing guide RNAs targeting GORAB were created in HEK293 Lenti-X cells (Takara 31966021) plated on 10 cm dishes the day before transfection. For each dish, 7 μg of each lentiU6-mGORAB gRNA plasmid alone (abm 224111140595; gRNA1 target 1-9 GGATTGGGCGGGCTTCTCTG; gRNA2 target 2-76 TTCACAGGAATTCGACGCTG; gRNA3 target 3-284 AGGTAGCTTTCCATCGCCAG), or all three in combination (2 μg of each gRNA plasmid), 3 μg of psPAX2 packing plasmid and 2 μg of pM2G envelope plasmid were transfected into HEK293 Lenti-X cells using 30 μL of Fugene-HD (Promega) in antibiotic-free DMEM. The medium was replaced the day after and the virus-containing medium was collected the following day, filtered through a 0.44 μm syringe filter unit and 5 ml were used for transduction of wild-type or vimentin KO MEFs with 1:1 ratio of medium containing virus to regular medium. The following day the medium was replaced with fresh medium containing puromycin (2 μg/ml), and cells were selected by culturing in medium containing antibiotic for 7 days. GORAB KO cells were diluted to single cells in 24 well plates. The GORAB KO data shown in the paper was generated with a clone generated using gRNA1. Similar results were obtained using gRNA3. The Vim GORAB double KO (dKO) cells were not able to proliferate from single cells and thus the dKO cell were used as a mixed population of cells obtained from the initial transduction.

### Proximity-dependent biotinylation

The proximity biotinylation method was adapted from Roux *et al*. (Roux et al., 2012). HeLaM or human skin fibroblast cells stably expressing BIOID-GORAB were incubated for 24 h in DMEM supplemented with 10% HyClone FBS, 1 mM L-glutamine, penicillin-streptomycin mix and 50 μM biotin. After three PBS washes, 4 × 10^7^ cells were lysed at 25°C in 1 mL lysis buffer (50 mM Tris-Cl pH 7.4, 0.5 M NaCl, 0.4% SDS, 5 mM EDTA, 1 mM DTT, and protease inhibitor cocktail (Calbiochem)) and sonicated (Bioruptor, Diagenode, Belgium) following the manufacturer’s instructions. Triton X-100 was added to 2% final concentration. After further sonication, an equal volume of cooled (4°C) 50 mM Tris-Cl pH 7.4 was added before additional sonication (subsequent steps at 4°C) and centrifugation at 14 000 rpm. Supernatants were incubated with 500 μL pre-equilibrated streptavidin-Dynabeads (MyOne Steptavadin C1, ThermoFisher) overnight. Beads were collected and washed twice for 8 min at 25°C (all subsequent steps at 25°C) in 1 mL wash buffer 1 (2% SDS in dH2O). This step was repeated once with wash buffer 2 (50 mM HEPES pH 7.5, 0.1% deoxycholate, 1% Triton X-100, 500 mM NaCl and 1 mM EDTA), once with wash buffer 3 (10 mM Tris-Cl pH 8.1, 250 mM LiCl, 0.5% NP-40, 0.5% deoxycholate and 1 mM EDTA) and twice with wash buffer 4 (50 mM Tris-Cl pH 7.4, and 50 mM NaCl). Samples to be analysed by mass spectrometry (MS) were washed twice in 50 mM NH_4_HCO_3_. On-bead tryptic digests were analysed by 1D Liquid chromatography–mass spectrometry (LC/MS/MS) by Sanford-Burnham Proteomic Facility (La Jolla, CA) using the following procedure. 4 μL of Tris(2-carboxyethyl)phosphine (TCEP) was added to 200 μl of beads/50 mM ammonium bicarbonate suspension mix and proteins were reduced at 40°C for 30 min. Iodoacetamide was added to 20 mM and proteins were alkylated at 30 min at RT in the dark. Mass spectrometry grade trypsin was added (1:20 ratio) for overnight digestion at 37°C. After digestion, magnetic beads were removed by centrifugation. Formic acid was added to the peptide solution to 2%, followed by desalting and on-line analysis of peptides by high-resolution, high-accuracy LC-MS/MS, consisting of a Michrom HPLC, a 15-cm Michrom Magic C18 column, a low-flow ADVANCED Michrom MS source, and a LTQ-Orbitrap XL (Thermo Fisher Scientific, Waltham, MA). A 120-min gradient of 10–30%B (0.1% formic acid, 100% acetonitrile) was used to separate the peptides. The total liquid chromatography time was 141 min. The LTQ-Orbitrap XL was set to scan precursors in the Orbitrap followed by data-dependent MS/MS of the top four precursors. Raw LC-MS/MS data were submitted to Sorcerer Enterprise (Sage-N Research, Milpitas, CA) for protein identification against the ipi.HUMAN.vs3.73 protein database, which contains semi-tryptic peptide sequences with the allowance of up to two missed cleavages. A molecular mass of 57 Da was added to all cysteines to account for carboxyamidomethylation. Differential search included 16 Da for methionine oxidation, and 226 Da on N terminus and lysine for biotinylation. Search results were sorted, filtered, statically analysed, and displayed using PeptideProphet and ProteinProphet (Institute for Systems Biology, Seattle, WA) (Nesvizhskii et al., 2003). The minimum trans-proteomic pipeline probability score for proteins was set to 0.95, to assure trans-proteomic pipeline error rate of lower than 0.01. The relative abundance of each identified proteins in different samples were analysed by QTools, an open source in-house developed tool for automated differential peptide/protein spectral count analysis (Brill et al., 2009).

### Preparation of cell lysates

MEFs were washed three times in PBS and lysed in ice-cold lysis buffer (50 mM Tris-HCl, pH 7.4, 150 mM NaCl, 1% Triton X-100, 1% protease inhibitor cocktail; 200 μl/well in 6 well plate) and incubated on ice for 30 min. The lysate was cleared by centrifugation at 16,000xg for 10 min at 4°C and the protein concentration was measured with the Bradford method.

### Immunoblotting

40 to 80 μg of cell lysate was loaded onto a 10% SDS-PAGE, transferred onto a nitrocellulose membrane followed by blocking in 5% non-fat dried milk in PBS supplemented with 0.15% Tween20 for 1 h at RT°, overnight incubation with primary antibodies at 4°C. HRP-conjugated antibodies were used as secondary antibodies with 1 h incubation at RT°. Protein bands were visualized using SuperSignal West Femto substrate (Thermo) on a Chemidoc imager (Bio-Rad).

### Immunofluorescence microscopy

Cells were fixed in3.7% paraformaldehyde (PFA) in PBS for 20 min at 37C. Then, cells were washed 2 times with PBS, quenched with 1 M Glycine and permeabilized by incubating the cells in 0.25% Triton X-100 in PBS for 5 min at RT. After three washes with PBS the cells were incubated 1 h in blocking solution (0.2% saponin, 1% FBS, 0.5% BSA in PBS) followed by primary antibodies incubation at RT for 1 h and 30 min. Coverslips were then washed three times in PBS for 5 min and incubated with the secondary antibodies supplemented with DAPI (200 ng/ml) for 1 h at RT, washed three times with PBS for 5 min, dried and mounted on slides using Prolong mounting solution. Images were acquired using an Olympus BX60 upright microscope equipped with a MicroMax cooled, slow-scan CCD camera (Princeton Instruments, Acton, MA) driven by Metaview software (University Imaging Corporation, West Chester, PA). Images were processed using ImageJ software (MacBiophotonics, Bethesda, MD). For the confocal and STED microscopy images were collected on a Leica TCS SP8 AOBS inverted gSTED microscope with the following confocal setting: pinhole 1 Airy unit, scan speed 400 Hz unidirectional and format 2048 × 2048; Alexa 488: 498–542 nm; Alexa-549: 564–619 nm; Alexa-647:646–713 nm using the 490 nm, 555 nm and 635 nm excitation laser lines and 592 nm, 660 nm and 775 nm depletion laser lines, respectively. STED images were deconvolved using Huygens Professional (Scientific Volume Imaging).

### Drug treatments

Medium containing BFA at a final concentration of 5 μg/ml was added to MEFs grown on coverslips, Cells were incubated for 5, 10, 20, 30 and 60 min in the presence of BFA, prior to fixation in 4% PFA. For BFA washout experiments, MEFs were treated for 90 min with BFA, washed three times with fresh medium and incubated for and additional 15, 30, 45, 60 and 90 min prior to fixation. For depolymerisation of microtubules, cold medium containing nocodazole at a final concentration of 8 μg/ml was added to MEFs and cells were cultured at 37°C for 2 h before fixation.

### EM Analysis of Golgi ultrastructure

MEF cells were grown on a 10 cm dish until they reached confluency. The samples were fixed by adding double concentrated fixative directly to the same amount of medium in the culture dish, so that the final fixative was 4% formaldehyde + 2.5% glutaraldehyde in 0.1 M HEPES buffer (pH 7.2). Then samples were post-fixed with reduced osmium (1% osmium tetroxide + 1.5% potassium ferrocynaide) in 0.1 M cacodylate buffer(pH 7.2) for 1 h, then in 1% uranyl acetate in water overnight. The samples were dehydrated in ethanol series infiltrated with TAAB LV resin and polymerized for 24 h at 60°C. Sections were cut with Reichert Ultracut ultramicrotome and observed with FEI Tecnai 12 Biotwin microscope at 100kV accelerating voltage. Images were taken with Gatan Orius SC1000 CCD camera.

### Image analysis and statistical tests

Colocalization, Golgi elements and total cell fluorescence were quantified using ImageJ/Fiji software. Golgi element number was determined by counting the structures/cells above intensity fluorescence threshold that had been selected based on WT MEF cells. Total intensity fluorescence was normalized for cell area. Statistical analyses were conducted using GraphPad Prism software (GraphPad Software, La Jolla, CA). The D’Agostino–Pearson and Shapiro–Wilk tests were used for comparison of the distribution of data with a Gaussian distribution. Depending on the result, an unpaired t-test or Mann–Whitney test was performed. In the case of an unpaired t-test, equality of variances between two groups was tested with an F-test. For multiple group comparisons, a one-way or two-way ANOVA followed by Dunnett’s test was performed. Statistical significance cut-offs were set as follows: * – p≤0.05, ** – p<0.01, *** – p<0.001 and **** – p<0.0001.

## Acknowledgements

We would like to thank Dr. Sara Koester (University of Gottingen) for providing the vimentin KO MEFs and May Tassbehji and Viki Allan (University of Manchester) for providing the GO patient cells and SW13 cells respectively. We are grateful to the FBMH Bioimaging core facility for their help with the microscopy and acknowledge the FBMH Electron Microscopy core facility (RRID: SCR_021147). This work was supported by a BBSRC project grant (ML and TV; BB/T000945/1), a BBSRC sLoLa award (ML, JS, MC; BB/T001984/1) and an MRC project grant (ML, TW; MR/N000366/1).

## Notes

### Competing Interest Statement

The authors have declared no competing interest.

